# TRPV5, TRPV6, TRPM6, and TRPM7 do not contribute to hair-cell mechanotransduction

**DOI:** 10.1101/204172

**Authors:** Clive P. Morgan, Hongyu Zhao, Meredith LeMasurier, Wei Xiong, Bifeng Pan, Matthew R. Avenarius, Michael Bateschell, Ruby Larisch, Anthony J. Ricci, Ulrich Müller, Peter G. Barr-Gillespie

**Affiliations:** Oregon Hearing Research Center & Vollum Institute, Oregon Health & Science University, Portland, OR 97239, USA; Department of Neuroscience, Scripps Research Institute, La Jolla, CA 92037; Department of Otolaryngology, Stanford University, Stanford, CA, 94305

## Abstract

The hair-cell mechanotransduction channel remains unidentified. We tested whether four transient receptor channel (TRP) family members, TRPV5, TRPV6, TRPM6, and TRPM7, participated in transduction. Using cysteine-substitution mouse knock-ins and methanethiosulfonate reagents selective for those alleles, we found that inhibition of TRPV5 or TRPV6 had no effect on transduction in mouse cochlear hair cells. TRPM6 and TRPM7 each interacted with the tip-link component PCDH15 in cultured eukaryotic cells, which suggested they could participate in transduction. Cochlear hair cell transduction was insensitive to shRNA knockdown of *Trpm6* or *Trpm7*, however, and was not affected by manipulations of Mg^2+^, which normally perturbs TRPM6 and TRPM7. To definitively examine the role of these two channels in transduction, we showed that deletion of either or both of their genes selectively in hair cells had no effect on auditory function. We suggest that TRPV5, TRPV6, TRPM6, and TRPM7 are unlikely to be the pore-forming subunit of the hair-cell transduction channel.

## Introduction

A central mystery in auditory neuroscience is the identity of the molecules making up the pore of the hair cell’s transduction channel, a cation-selective channel that responds to mechanical stimuli and produces a receptor potential in the cell. Transduction channels are gated by tension in tip links, thin extracellular filaments made from the cadherins CDH23 and PCDH15 (Siemens et al., 2004; Sollner et al., 2004; Alagramam et al., 2011). PCDH15 is located at the base of the tip link (Kazmierczak et al., 2007), where the transduction channel is located (Beurg et al., 2009); indeed, PCDH15 may bind directly to the transduction channel (Powers et al., 2012). The ion channel HCN1 has been reported to bind to PCDH15 (Ramakrishnan et al., 2009; Ramakrishnan et al., 2012), but *Hcn1* knockouts have normal hearing (Horwitz et al., 2010). By contrast, strong evidence suggests that the channel complex contains the transmembrane proteins TMC1 and TMC2 (Kawashima et al., 2011; Pan et al., 2013), TMIE (Zhao et al., 2014), and LHFPL5 (Xiong et al., 2012); moreover, PCDH15 interacts with both TMC1 and TMC2 (Maeda et al., 2014; Beurg et al., 2015). Nevertheless, evidence that any of these proteins contributes directly to the ion-permeation pathway is lacking.

Because they carry out a wide variety of cellular functions, including sensory transduction, the 33-member transient receptor potential (TRP) family might well contain the transduction-channel pore molecule (Venkatachalam and Montell, 2007; Zanini and Göpfert, 2014). Many TRP channels are present in the inner ear; a comprehensive quantitative RT-PCR investigation of TRP channel expression during cochlear development detected transcripts for 30 different TRP genes (Asai et al., 2010). Several TRP channels have been advanced specifically as candidates for the transduction channel, but all evidence so far has ruled out these candidates (Fettiplace and Kim, 2014).

Our preliminary evidence raised the possibility that TRPV5, TRPV6, TRPM6, or TRPM7 might be part of the transduction channel. TRPV channels play key roles in transduction in fly mechanoreceptors (Gong et al., 2004; Kernan, 2007; Lehnert et al., 2013; Zhang et al., 2013). As they are highly selective for Ca^2+^ (Owsianik et al., 2006), TRPV5 and TRPV6 are less likely as candidates for the transduction channel than other channels; nevertheless, the transduction channel’s permeability and conductance can change substantially depending on other components expressed (Xiong et al., 2012; Beurg et al., 2015). TRPM7, a member of the melanostatin subfamily, has been implicated in mechanosensation (Bessac and Fleig, 2007; Numata et al., 2007a; Numata et al., 2007b; Wei et al., 2009; Xiao et al., 2015). TRPM7 and its closely related Paralog TRPM6 uniquely possess a C-terminal kinase domain; these channels are thought to regulate Mg^2+^ homeostasis as well as cell death, proliferation, differentiation, and migration (Chubanov and Gudermann, 2014; Fleig and Chubanov, 2014). *Trpv5, Trpv6, Trpm6*, and *Trpm7*transcripts were all detected in cochlea (Asai et al., 2010); *Trpm6* levels increased during development, *Trpv6* and *Trpm7* decreased, and *Trpv5* was seen in only a fraction of the samples. Both *Trpm6* and *Trpm7* were detected in a cochlear cDNA library using conventional PCR, but *Trpv5* and *Trpv6* were not (Cuajungco et al., 2007).

Because the cysteine-substitution mutations in TRPV5 (S556C) and TRPV6 (M527C) render these channels sensitive to methanethiosulfonate (MTS) reagents, we generated S556C-*Trpv5* and M527C-*Trpv6* knock-in mice. Mechanotransduction was insensitive to MTS reagents, however, in both heterozygote and homozygote hair cells from these lines. Similarly, mechanotransduction in hair cells was not affected by conditions that should alter TRPM6 or TRPM7 function, and conditional deletion of the *Trpm6* and *Trpm7* genes in hair cells did not affect auditory function. We suggest that TRPV5, TRPV6, TRPM6, and TRPM7 should be added to the long list of ion channels that have been shown to not be part of the transduction channel (Horwitz et al., 2010).

## Materials and Methods

### Animal research

This study was carried out in compliance with the Animal Welfare Act regulations and the Office of Laboratory Animal Welfare—Public Health Service Policy on Humane Care and Use of Laboratory Animals. Protocols were approved by the Institutional Animal Care and Use Committees (IACUC) of Oregon Health & Science University, Stanford University, and Scripps Research Institute.

### Generation of mouse lines

For *M527-Trpv5*, we generated a targeted construct that contained (5’ to 3’) a short homology arm (1.8 kb) with an added loxP-flanked neomycin resistance cassette, exon 13 including the codon encoding M527 to a cysteine and an introduced *ScaI* site for screening, a long homology arm (5.3 kb), and a diphtheria toxin cassette. The targeting construct was linearized with *XhoI* for electroporation into ES cells. Screening was carried out on the 5’ and 3’ arms using PCR, with TOPO-cloning of PCR products to verify that the correct genomic DNA region was targeted. Mice were generated from the correctly-targeted ES cells by the University of Cincinnati Gene-Targeted Mouse Service. After blastocyst injections, founders were identified by coat color chimerism and were bred to C57BL/6 mice for >6 generations. Mice were genotyped using PCR, with AAATGGGAACCAGATTCATCTCA as the forward primer and ACTATACAAAAGGGTAACCTACCCACA as the reverse primer. The amplified samples were digested with *BamHI* (the knock-in removes a *BamHI* site) and analyzed by electrophoresis.

For *S556C-Trpv6*, we generated a targeted construct that contained (5’ to 3’) a diphtheria toxin cassette, a long homology arm (5.2 kb), exon 13 including the codon encoding the S556C alteration and an introduced *AvrII* site for screening, and a short homology arm (1.65 kb) with an added loxP-flanked neomycin resistance cassette. Screening of targeted ES cells was carried out using Southern blotting with 5’ and 3’ probes, and targeted mice were generated, screened, and bred as above. Mice were genotyped using PCR, with CCAGTGTCTCATGCTTATATCC as the forward primer and TAGATGTCATGAAGATTAAAGG as the reverse primer. The amplified samples were digested with *ScaI* (the knock-in adds a *ScaI* site) and analyzed by electrophoresis.

We obtained *Trpm6*^*tm1a(KOMP)Wtsi*^ ES cells from the UC Davis KOMP repository (clones EPD0741_2_G10 and EPD0741_2_G11). After blastocyst injections, founders were identified by coat color chimerism and targeting was verified by PCR analysis. Founders were bred to C57BL/6, and the FRT-neo cassette was removed by crossing with the *Flp* deleter line B6;SJL-Tg(ACTFLPe)9205Dym/J (Jackson Laboratories). PCR was used to verify loss of the cassette. These mice were referred to as *Trpm6*^*fl*^. To produce *Trpm6*^*CKO*^ mice, we used the *Atoh1-Cre* mouse line (Matei et al., 2005), which recombines floxed genes in hair cells with >99% efficiency (Avenarius et al., 2017). Floxed mice were genotyped using PCR, with GCTCCTCAGGGTTCCTCCAGTCTGT as the forward primer and GCAAGGACAAGAGGGCGTCAGAGC as the reverse primer. The amplified samples were analyzed by electrophoresis; a wild-type allele produces a band of 670 bp, while the floxed-allele band is 833 bp.

We generated a mouse allele with the putative ion-conductance pore of *Trpm7* flanked by *loxP* sites. The 5’ arm (4948 bp) was PCR-amplified using a forward primer that introduced an *ApaI* site (GCGGGGCCCTGGGTGATTGACATTTCATTCCAAGT) and a reverse primer that introduced a *SaII* site (GCGGTCGACTGTCAACTAGCAATGGAAATGCAGACTT). This fragment was introduced into the pBS-FRT-Neo-FRT plasmid. The 3’ arm (2987 bp) was PCR-amplified using the forward primer GCGCTCGAGGTGTATATAAGAATTGTCTCAGGATAGT and reverse primer GCGCCGCGGCCTCTTATCCTGTTTCTCTACATGTGT. A separate middle piece was prepared as a *SalI/NotI* fragment flanked by loxP sites; it was PCR-amplified using the forward primer GCGGTCGACGGTTTTGCCTTATATTTGCAAGGCATA and reverse primer GCGGCGGCCGCCCATTACCATCATTCCTTGAAGTGGCTTT. The middle piece was cloned into the 5’ arm vector at *SalI* and *NotI* sites; the 3’ arm was then cloned into this latter piece (5’ and middle). Founders were bred to C57BL/6, and the FRT-neo cassette was removed as above (producing *Trpm7*^*fl*^ mice). *Trpmi*^*CKO*^ mice were generated with *Atoh1-Cre* as above. Floxed mice were genotyped using PCR, with CCATACTGGATGATTTTTGGTGAAGTTTATGCA as the forward primer and CACAAACAAGGAAGGGAAGAGTTTTAATATCCA as the reverse primer. The amplified samples were analyzed by electrophoresis; a wild-type allele produces a band of 514 bp, while the floxed-allele band is 633 bp.

### Heterologous expression in HEK cells and immunoprecipitation

For coimmunoprecipitation assays, we used HEK293T cells grown in six-well tissue culture plates. Cells were transfected with Effectene transfection reagent (Qiagen, #301425) by preparing the DNA complex according to the manufacturer’s suggested protocol in PCR tubes, then adding the DNA complex to the cells. Cells were harvested at 48 hr, centrifuged briefly (16,000g for 5 sec), and the supernatant removed. To the pellet, 300 μl of cold lysis buffer containing PBS, 1% Triton X-100, 0.5% NP-40, and protease inhibitor cocktail (Sigma, #P8340) was added, and the cells were then sonicated on ice using a tip sonicator (Ultracell Sonicator, Sonics) set at 25% power. The lysate was then centrifuged at 16,000 g for 20 min at 4°C, and the supernatant was separated from the pellet. A total of 10 μl of HA agarose (Sigma, #A2095, 20 μl slurry) was added and the mixture was rotated overnight at 4°C. The immunoprecipitate mixture was centrifuged at 16,000 g for 1 sec, then washed 2x with 300 μl lysis buffer and 1X with PBS. After the washes, 32 μl of 2x SDS sample buffer (Invitrogen) was added to the agarose beads, and the mixture was incubated for 10 min at 70°C. The agarose beads were removed using a spin filter (Costar #8163), and 5 μl of 10x reducing agent (Invitrogen, #NP0007) was added to the protein solution. The mixture was again incubated at 70°C for 10 min. The protein samples were separated on a 3-8% Tris-acetate gel (Invitrogen, #EA03755BOX) for 30-40 min. Proteins in the gel were then transferred to PVDF membrane (Millipore, #IPVH00010) using a semi-dry setup (Bio-Rad, Transblot SD). Towbin buffer (25 mM Tris, 192 mM glycine, pH 8.3) with 10% MeOH and without SDS were used as transfer buffer. After transferring for 40-50 min, the blot was washed 1x in PBS and blocked in blocking buffer (PBS containing 10% FBS, 0.1% Tween) for 30 min, during which primary antibody were diluted in blocking buffer (1:1000) and incubated at room temperature for 10-20 min. The blot was incubated with primary antibody for 2 hr at room temperature. After washing the blot, HRP-coupled secondary antibody, diluted 1:100 in blocking buffer, was incubated with the blot for 30 min. After washing, SuperSignal West Pico Chemiluminescent Substrate (Pierce, #34077) was used for detection.

Anti-HA agarose or anti-V5 agarose (Sigma, #A7345) were used to pull down the HA-tagged TRP channels. In the case of PCDH15 pull down, we used 10 μg of PB811, a rabbit antibody against PCDH15 (Kazmierczak et al., 2007), or 10 μg of anti-HRP antibody (Jackson Immunoresearch) with 10 μl of Ultralink immobilized protein A (Pierce, #53139). For protein immunoblotting, PCDH15 was detected with PB811, TRP channels were detected with monoclonal anti-HA antibody (Applied Biological Materials, #G036).

### Mouse hair-cell mechanotransduction: *Trpv5* and *Trpv6* experiments

For tissue preparation, organ of Corti explants from the mid apex region were dissected from mice of either sex at ages P6-P9. The tissue was placed into the recording chamber and held in place with single strands of dental floss. A bath perfusion and an apical perfusion pipette (50 μm diameter) was placed into the chamber and external solution was perfused over the tissue. The external solution contained (in mM) 140 NaCl, 2 KCl, 2 CaCl_2_, 2 MgCl_2_, 10 (4-(2-hydroxyethyl)-1-piperazineethanesulfonic acid) (HEPES), 6 glucose, 2 pyruvate, and 2 creatine, and was balanced to a pH of 7.4 and osmolality of 300-310 osmol/l. After turning off apical perfusion for up to 30 minutes, inhibitors (MTSET or MTSES) were applied by hand while recording mechanotransduction (MET) currents. Inhibitors were aliquoted into single use samples and kept frozen until a recording was obtained. The inhibitor was put into solution and rapidly applied to the recording chamber within 1 min of dissolving to avoid any breakdown of the compound. Bath flow was turned on after 30 minutes of recording from a given cell. Only one cell was recorded per mouse and recordings were done blindly to animal genotype.

Electrophysiological recordings from hair cells were recorded as previously described (Peng et al., 2016). Thick-walled borosilicate patch pipettes, (2.5-3.5 MΩ) were used to establish whole-cell recordings from inner hair cells. The internal solution contained (in mM) 125 CsCl, 10 HEPES, 1 1,2-bis(o-aminophenoxy)ethane-N,N,N′,N′-tetraacetic acid (BAPTA), 5 ATP, 5 creatine phosphate, and 3.5 MgCl_2_; the pH was adjusted to 7.0 and osmolality 290-295 osmol/l. An Axopatch 200b amplifier was used to control electrode voltage and monitor current. The amplifier was controlled via JClamp software coupled to an IOtech interface. Recordings were included for data analysis if MET current amplitudes were stable prior to drug application and if the leak currents were less than 100 pA. For 70 cells recorded, the uncompensated series resistance was 14.6 ± 7 MΩ and capacitance was 5.8 ± 0.8 pF. Data were included for analysis only if the series resistance changed less than 10% during recording.

Hair bundles were stimulated in either of two ways (Peng et al., 2016). A glass probe affixed to a piezo stack controlled by JClamp software was placed in the central region of the hair bundle. The tip was shaped to that of the angled outer hair cell bundle. Driving voltage was filtered at 3 kHz to limit piezo resonance. Hair bundles were also stimulated with fluid motion driven by a Picospritzer.

### Mouse hair-cell mechanotransduction: *Trpm6* and *Trpm7* experiments

Whole cell recording were carried out on outer hair cells (OHCs) of either acutely isolated or cultured cochleae. Transducer currents were sampled at 100 KHz with a patch-clamp amplifier operated by Patchmaster 2.35 software (EPC10-USB, HEKA). The patch pipette was filled with an intracellular solution containing (in mM) 140 KCl, 1 MgCl_2_, 0.1 EGTA, 2 Mg-ATP, 0.3 Na-GTP and 10 H-HEPES, pH 7.2). For the magnesium test, MgCl_2_ was raised to 3 mM (3 Mg-SI) or removed (0 Mg-SI). The artificial perilymph contained (in mM): 144 NaCl, 0.7 NaH_2_PO4, 5.8 KCl, 1.3 CaCl_2_, 0.9 MgCl_2_, 5.6 glucose, and 10 H-HEPES, pH 7.4. OHCs were clamped at −70 mV. Borosilicate glass with a filament (Sutter) was pulled with a pipette puller (P-97, Sutter), and polished to resistances of 3-5 MΩ with a microforge (MF-830, Narishige). Hair bundles were deflected with a glass pipette mounted on a piezoelectric stack actuator (P-885, Physik Instrument). The tip of the pipette was fire-polished to 4-6 μm in diameter to fit the shape of OHC bundles. The actuator was driven with voltage steps that were low-pass filtered at 10 KHz with an eight-pole Bessel filter (900CTF, Frequency Devices).

### Gene knockdown of cochlear hair cells

OHCs were injectoporated with shRNAs against *Trpm7* transcripts as previously described (Xiong et al., 2014). In brief, the organ of Corti from P1 mice was cut into 3 pieces, then placed in DMEM/F12 medium containing 10% FBS and 1.5 μg/ml ampicillin. For electroporation, a glass electrode (2 μm diameter) was used to inject shRNA plasmid (pSicoR system, 0.5 μg/μl in 1x HBSS) into the cochlea, between the hair cells. A series of 3 pulses was applied at 1 sec intervals with a magnitude of 60V and a duration of 15 msec by an electroporator (ECM 830, BTX). The tissues were cultured for 3 days in vitro (DIV). Before electrophysiological recording, the dish with organotypic cultures was changed to artificial perilymph. OHCs with shRNA expression were selected by GFP signal.

### Other methods

Gene-gun transfection and visualization of fluorescence protein-tagged constructs was carried using an optimized procedure (Zhao et al., 2012). For biolistic transfection, mouse cochleas were dissected at P6, subjected to gene-gun transfection, and then cultured for 1 day. Surface biotinylation was carried out with NHS-LC-Sulfo-biotin (Elia, 2008). Shotgun mass spectrometry on immunoprecipitates was carried out using a Thermo Velos ion-trap mass spectrometer; peptides and proteins were identified using the PAW pipeline and SEQUEST searches (Wilmarth et al., 2009; Krey et al., 2014). ABRs were carried out as described (Schwander et al., 2007; Ebrahim et al., 2016).

## Results

### Evidence for role of TRPV5 and TRPV6 in transduction

While localization of PCDH15 to stereocilia tips does not depend on USH1C (Lefevre et al., 2008), PCDH15 and USH1C can bind directly under some conditions (Adato et al., 2005; Reiners et al., 2005), suggesting that USH1C might couple the transduction channel to PCDH15 under some conditions. Sequences of 33 TRP channel genes were scanned for PDZ binding interfaces (PBIs); seven that had candidate PBIs were tested in yeast two-hybrid assay with USH1C as the bait (Fig. 1A). Of these, only TRPV6 interacted with USH1C (Fig. 1B). We carried out RT-PCR for *Trpv6* transcripts and those for *Trpv5*, its close homolog, and found robust expression in vestibular tissues (Fig. 1C). Although immunocytochemistry experiments did not reliably detect either TRPV5 or TRPV6, the low levels of ion channels like TRP channels make them very difficult to detect. Together our data raised the possibility that TRPV6 interacts with USH1C in hair cells, thus making TRPV6 (and closely related TRPV5) a plausible transduction-channel candidate.

**Figure 1.**
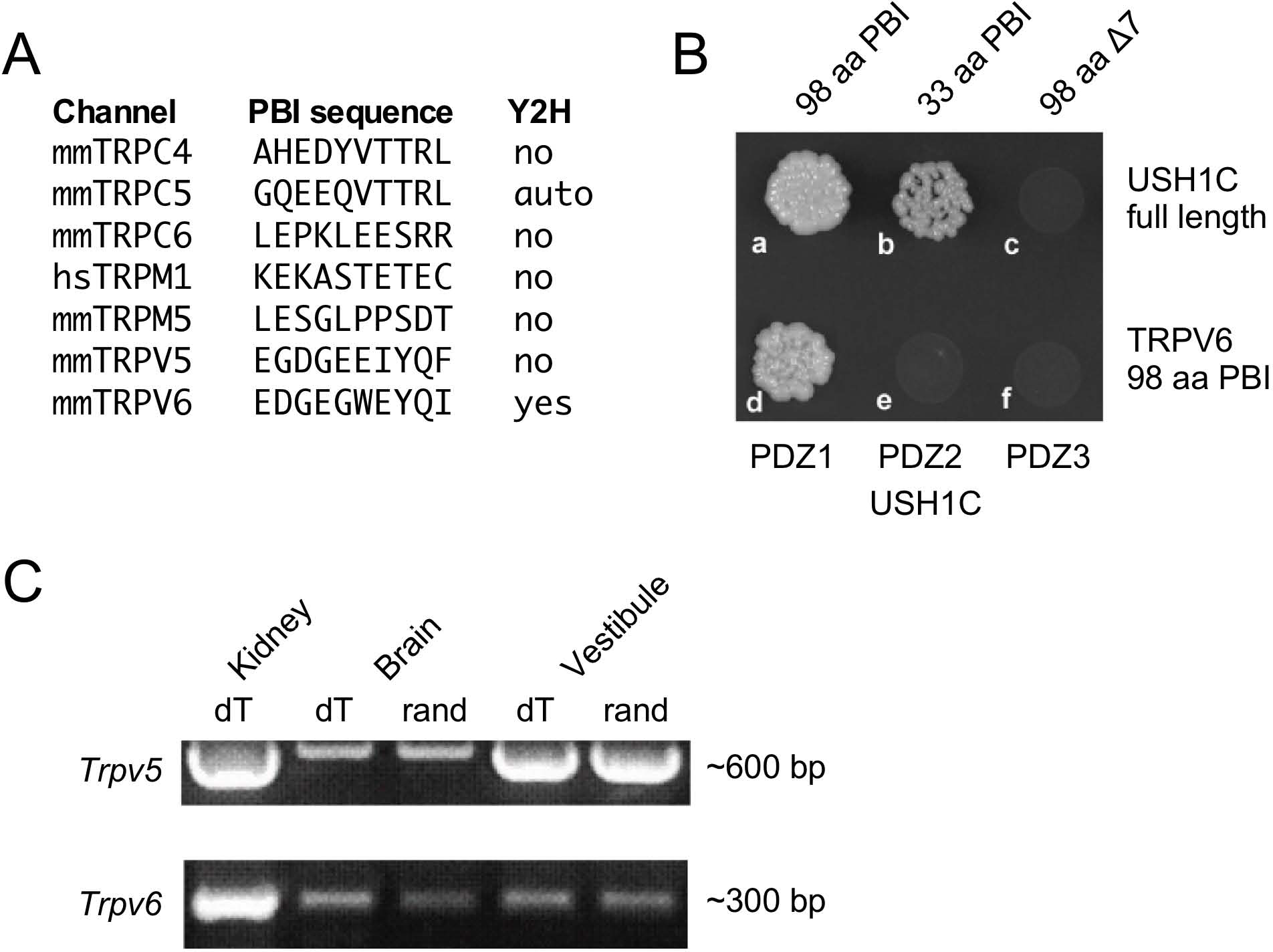
Characterization of TRPV5 and TRPV6. ***A***, Seven TRP channels (of 33) with candidate PDZ-binding interface motifs. Yeast two-hybrid (Y2H) results examining interactions with USH1C (harmonin) are indicated. ***B***, The 33 terminal amino acids of the TRPV6 PBI interact with USH1C, but deletion of the C-terminal seven amino acids abolishes binding (top row). The interaction of TRPV6 is through PDZ1 of USH1C (bottom row). ***C***, RT-PCR (with reverse transcription primed either with oligo dT or random hexamers) showing that mRNAs for TRPV5 and TRPV6 are present in vestibular RNA.

### MTS reagents do not affect transduction in *S556C-Trpv5* and M527C-*Trpv6* hair cells

Individual knockouts of *Trpv5* or *Trpv6* had no reported effects on auditory or vestibular function (Hoenderop et al., 2003; Bianco et al., 2006). These two genes are closely linked on human chromosome 7 or mouse chromosome 6 (Hoenderop and Bindels, 2008), so double knockouts cannot easily be created by breeding. To determine whether these channels participate in hair-cell transduction, we developed a method whereby inhibition of a single channel subunit should lead to a dominant-negative effect on the tetrameric channel (Voets et al., 2004a). Note that TRPV5 and TRPV6 can co-assemble into a single tetramer (Hellwig et al., 2005). We used cysteine-substitution mutagenesis to create *Trpv5* and *Trpv6* alleles that were sensitive to MTS reagents. Residue-scanning experiments with cysteine substitution and MTS inhibition demonstrated that the S556C mutation rendered TRPV5 sensitive to sulfhydryl reagents (Dodier et al., 2004); similar experiments with TRPV6 identified M527C as a suitable mutation that created an MTS-sensitive allele (Voets et al., 2004a). We used standard gene-targeting methods to introduce mutations in the mouse genome that produced S556C-TRPV5 and M527C-TRPV6 expression (Fig. 2).

**Figure 2.**
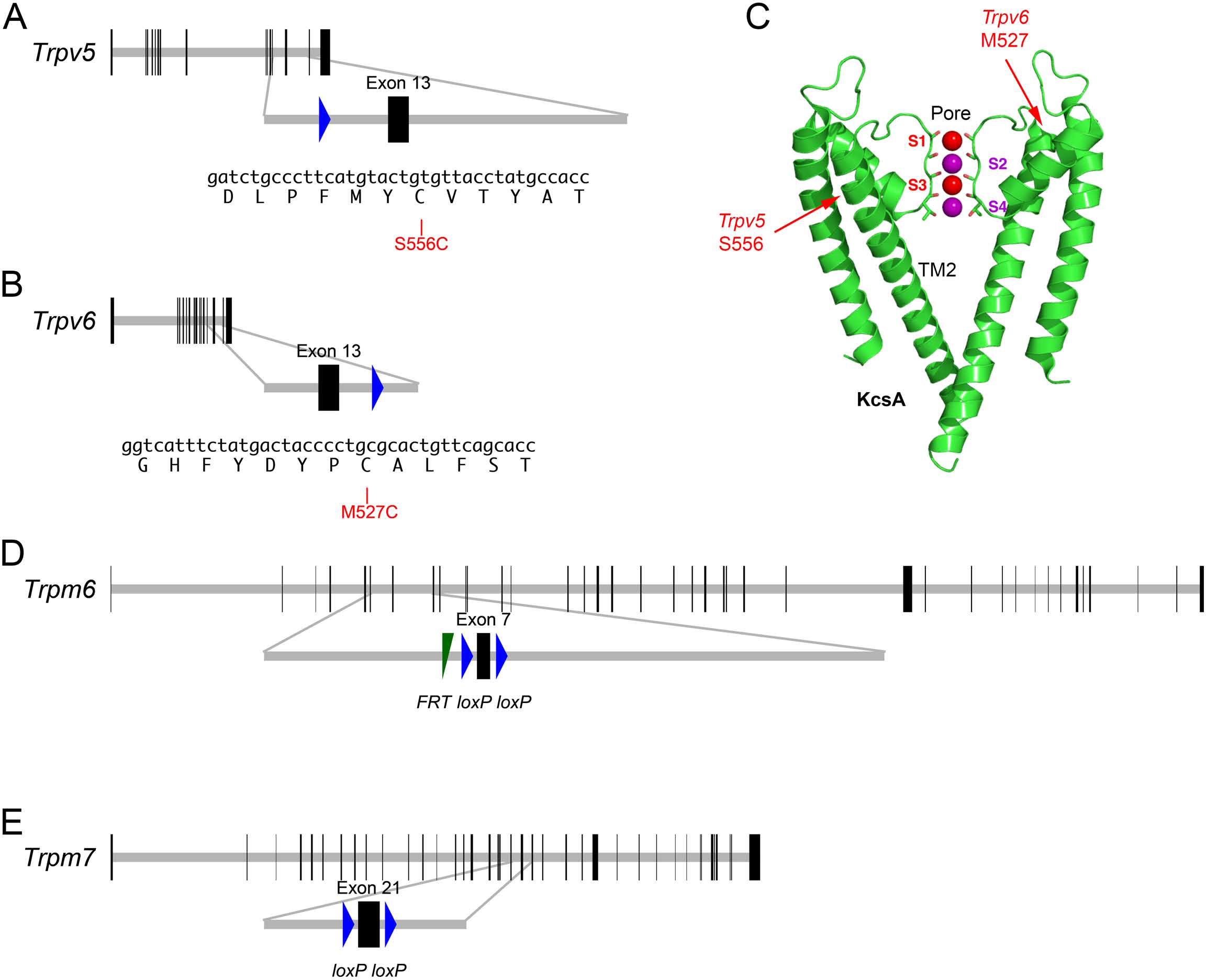
Targeted mutations in mouse *Trpv5*, *Trpv6*, *Trpm6*, and *Trpm7* genes. ***A***, Top, *Trpv5* gene structure. Horizontal gray line indicates scaled length of gene with coding exons, which are shown as vertical black rectangles. Middle, magnification of targeted exon. The *loxP* site remaining after excision of the *neo* cassette is shown in blue. Bottom, nucleotides and protein translation for region targeted. The S556C mutation is indicated. ***B***, M527C-Trpv6 targeting. ***C***, Structure of the KcsA ion channel (image from https://commons.wikimedia.org/wiki/File:1K4C.png), along with the locations of the residues analogous to S556C-*Trpv5* and M527C-*Trpv6*. The pore and transmembrane domain 2 (TM2) are indicated as well. ***D***, Structure of *Trpm6*^*fl*^ allele; exon 7 is flanked by *loxP* recombination sites; a residual *FRT* site remaining after excision of the *Flp* cassette is indicated. ***E***, Structure of *Trpm7*^*fl*^ allele; exon 21 is flanked by *loxP* sites.

S556C-*Trpv5* and *M527C*-*Trpv6* mice were viable and exhibited no apparent behavioral abnormalities as heterozygotes or homozygotes. Transduction currents of heterozygous S556C-*Trpv5* and *M527C*-*Trpv6* hair cells under control conditions appeared normal in their amplitudes, kinetics, and adaptation (Fig. 3A,C). In wild-type mice, MTS reagents like MTSET and MTSEA had no effect on mechanotransduction in the absence of the sensitizing *Trpv* alleles (Fig. 3), which demonstrates the suitability of the cysteine-substitution approach for probing the transduction channel.

**Figure 3.**
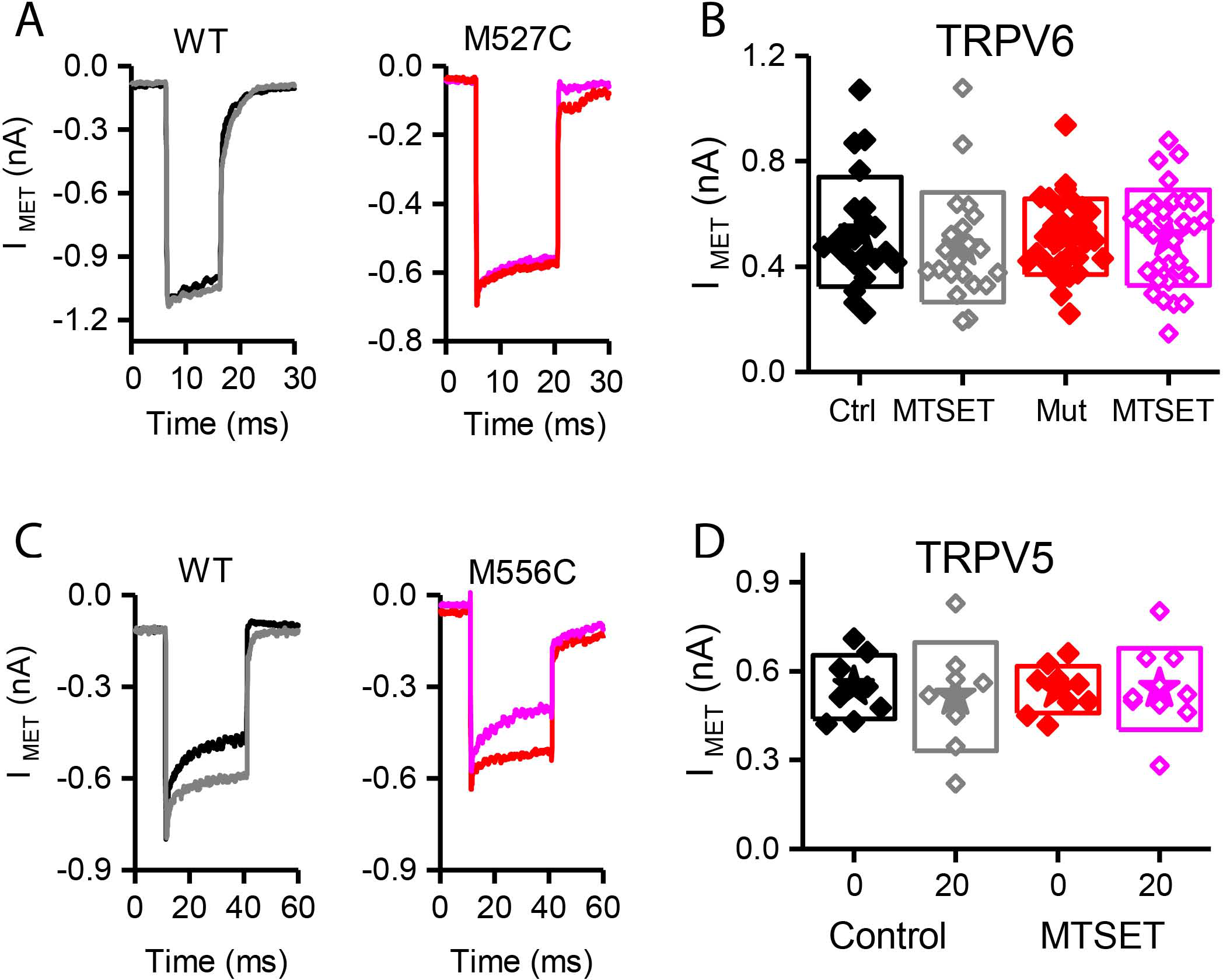
MTS reagents do not inhibit hair-cell transduction in M527C-*Trpv5* and S556C-*Trpv6* hair cells. ***A***, Hair-cell transduction currents from wild-type and M527C-Trpv5 heterozygous mice before and after treatment with 1 mM MTSET, applied in the extracellular solution. Color code is the same as in panel B. ***B***, Summarized data. Box plots show individual data points with the star indicating the mean and the box representing the standard deviation of the mean. Each point was obtained from one mouse and data were collected blinded to genotype. No significant differences were observed between any of the groups. ***C***, M527C-*Trpv5* wild-type and heterozygote hair cells before and after treatment with 1 mM MTSET, applied in the intracellular solution. Color code is the same as in panel D. ***D***, Summarized data for experiments similar to those in C; the x-axis represents time (in minutes) after obtaining the wholecell configuration.

Unfortunately, treatment of S556C-*Trpv5* or M527C-*Trpv6* hair cells with 1 mM MTSET or MTSES, which inhibit TRPV5 or TRPV6 activity by >95% in tissue culture cells (Dodier et al., 2004; Voets et al., 2004a), had no significant effects on mechanotransduction (Fig. 3). In the experiments illustrated in Fig. 3A, hair cells were treated with MTSET applied in the external solution; no changes in the maximal transduction current were detected under these conditions for either wild-type or M527C-*Trpv6* heterozygotes over the course of the recording (Fig. 3B). S556C-*Trpv5* heterozygote or homozygote hair cells were not inhibited by MTSET applied in the extracellular solution (data not shown). While the TRPV channels should be oriented in the membrane with the substituted residues exposed to the extracellular solution, there was a remote chance that the channels could be topologically flipped. We therefore also examined whether S556C-*Trpv5* hair cells could be inhibited with the MTS reagent applied through the recording electrode (Fig. 3C-D). There was no difference in transduction current amplitude between wild-type and S556C-*Trpv5* hair cells, however, even after 20 min after establishment of the whole-cell recording configuration (Fig. 3D).

### TRPM6 and TRPM7 co-immunoprecipitate with PCDH15 in tissue-culture cells

Based on its participation in mechanotransduction in other cell types, TRPM7 (and its close paralog TRPM6) is another plausible transduction channel candidate. We used heterologous expression of mouse proteins in HEK cells to investigate whether the TRPM channels can interact with PCDH15 (Fig. 4). We cloned full-length *Trpm6* and *Trpm7* from cDNA prepared from mouse inner ear, tagged them with the HA epitope tag, and co-expressed them with native, full-length PCDH15 in HEK293 cells. After solubilizing proteins and immunoprecipitating, when HA-tagged TRPMs were expressed, we could immunoprecipitate PCDH15 with anti-HA beads but not with negative-control anti-V5 beads (Fig. 4A). Likewise, immunoprecipitation with the anti-PCDH15 antibody PB811 selectively co-precipitated TRPM6 and TRPM7 (Fig. 4B). Co-expression of TRPM6 and TRPM7 consistently led to substantially more PCDH15 immunoprecipitated than with either alone (Fig. 4A).

**Figure 4.**
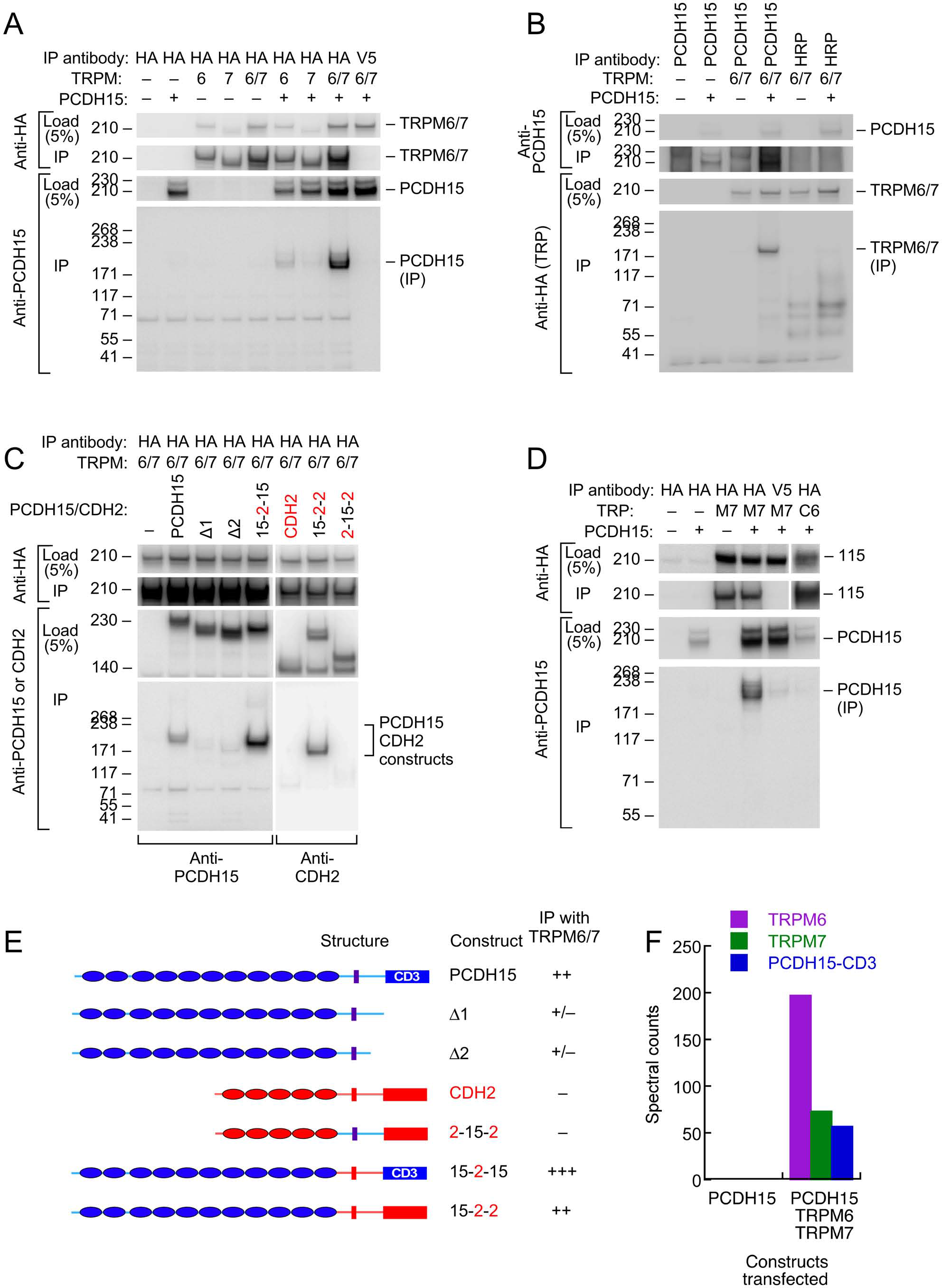
Interaction of PCDH15 with TRPM6 and TRPM7. ***A***, Lysates were immunoprecipitated with anti-HA agarose, and probed with anti-HA (top panels) or anti-PCDH15 antibody (bottom panels). PCDH15 was only immunoprecipitated by anti-HA if TRPM6 or RPM7 were present. No PCDH15 was precipitated by the control antibody (V5). ***B***, Lysates were immunoprecipitated with protein A/G agarose and either PB811 antibody (for PCDH15) or HRP antibody (control), then probed for TRP channels with anti-HA antibody. TRPM6 and TRPM7 were only precipitated with co-expressed with PCDH15 and precipitated with an anti-PCDH15 antibody. ***C***, Lysates were immunoprecipitated with anti-HA agarose, and probed with anti-PCDH15 antibody (left panels) or CDH2 antibody (right). Two C-terminal deletion proteins of PCDH15, Δ1 and Δ2, did not immunoprecipitate with TRPM6/7. Chimeras that have N-terminal extracellular domains of PCDH15 (15-2-15 and 15-2-2) immunoprecipitated with TRPM6/7 more strongly than PCDH15. The chimera that does not contain extracellular domains of PCDH15 (2-15-2) did not immunoprecipitate with TRPM6/7. ***D***, Lysates were immunoprecipitated with anti-HA agarose and probed with an anti-PCDH15 antibody (PB811). The last two lanes are two controls: anti-V5 agarose as a control for non-specific binding of the agarose with PCDH15, and TRPC6 as a TRP channel outside the TRPM family. Both controls have much lower levels of PCDH15 immunoprecipitated. ***E***, Summary of PCDH15 constructs (N- to C-terminal) used in panel C. Chimeras with CDH2 used the CD3 splice form of PCDH15. ***F***, Mass spectrometry spectral counts for TRPM6, TRPM7, and PCDH15 in large-scale immunoprecipitation experiments using ant-HA agarose. No PCDH15 was immunoprecipitated when it was expressed alone, but large amounts were co-precipitated with TRPM6 and TRPM7.

To demonstrate the selectivity of the interaction, we showed that a different cadherin, CDH2 (N-cadherin), was not immunoprecipitated with HA-tagged TRPM6 or TRPM7 (Fig. 4C), nor was PCDH15 immunoprecipitated with HA-TRPC6, a different TRP channel (Fig. 4D). Co-expression was important; if HA-tagged TRPMs were expressed in separate samples from those of PCDH15 and the cell extracts were mixed, no PCDH15 was immunoprecipitated with HA-tagged TRPMs (data not shown). Deletion of the C-terminus of PCDH15 eliminated the interaction, but chimeras of PCDH15 and CDH2 that contained only the N-terminal extracellular domain of PCDH15 robustly co-precipitated TRPM6 and TRPM7 (Fig. 4C).

To determine whether any endogenous HEK proteins mediated the interaction between the TRPMs and PCDH15, we carried out a larger scale immunoprecipitation and subjected the final eluates to tandem mass spectrometry. When expressed by itself and immunoprecipitated by HA-agarose, no PCDH15 was detected in eluates. By contrast, when TRPM6 and TRPM7 were co-expressed with PCDH15, all three proteins were detected at high levels in the immune pellet eluates (Fig. 4F). Besides human heat shock protein cognate 71, no additional proteins co-immunoprecipitated at apparent stoichiometric levels, suggesting that the TRPM-PCDH15 interaction is direct.

### tdTomato-TRPM7 localizes to hair cell lateral membranes at the cell apex

Localization of TRP channels by immunocytochemistry is generally unreliable (Gilliam and Wensel, 2011). Instead, we generated a *tdTomato-Trpm7* plasmid and transfected it into hair-cell progenitors at P6 using gene-gun transfection. Transfection efficiency was poor, presumably because of the large size of the expression construct. We were able to identify several transfected cells using co-expressed ZsGreen, and found tdTomato signal that was largely localized to the lateral membranes of hair cells, close to the adherens junctions (Fig. 5A). While not detected in stereocilia, the location of tdTomato-TRPM7 close to the hair cell’s apical domain raised the possibility that small amounts of TRPM7 could be transported to stereocilia membranes.

**Figure 5.**
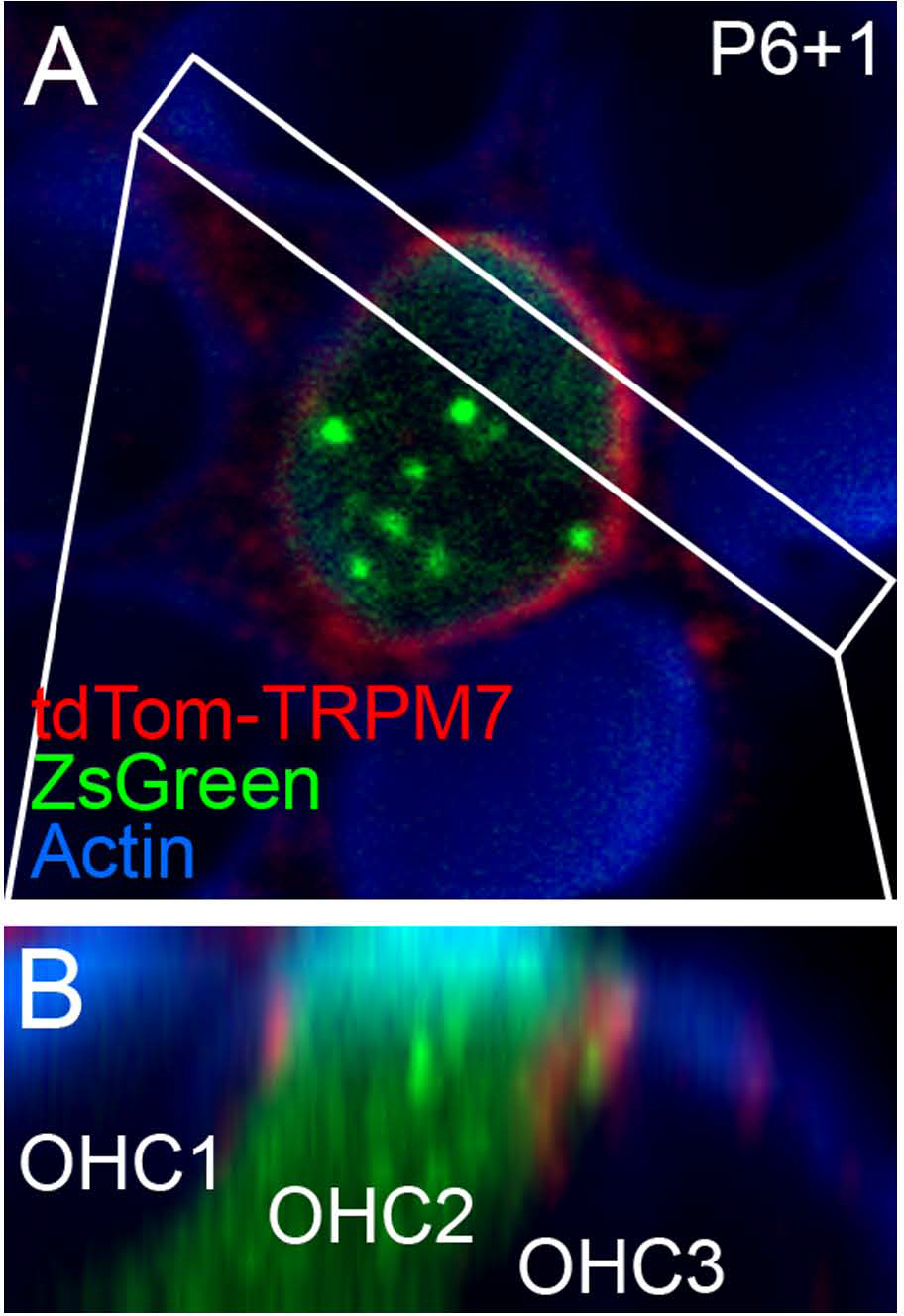
Localization of TRPM7 in the inner ear. ***A***, Single x-y slice through apical region of outer hair cells. One transfected cell, identified by ZsGreen signal, shows tdTomato signal at the lateral membrane. Box indicates region used for x-z reslice in B. P6+1, cochlea dissected at P6 and maintained in culture for 1 day. ***B***, Reslice of stack shows lateral view of transfected hair cell (in row 2; OHC2) and two other untransfected cells. tdTomato-TRPM7 signal was located near apex of cell.

### Identification of a novel *Trpm7* splice form in inner ear

When cloning full-length *Trpm6* and *Trpm7* from cDNA prepared from mouse inner ear, we also identified novel splice forms that skipped exon 20 of each gene, which in both genes encodes transmembrane helix 2 (TM2). Using RT-PCR analysis, we found that while the canonical *Trpm7* splice form was expressed in all mouse tissues, while *Dex27* (the *Trpm7* splice form lacking exon 20) was restricted to inner ear, heart, and liver. *Dex26* was expressed in a variety of tissues, including the inner ear.

Interestingly, both DEX26 and DEX27 were readily expressed in HEK293 cells and were present on the cell surface, as assessed by a surface biotinylation assay (Fig. 6A). DEX26 and DEX27 both co-immunoprecipitated PCDH15 as efficiently as the canonical splice forms (Fig. 6B), and could be co-immunoprecipitated with the canonical forms.

**Figure 6.**
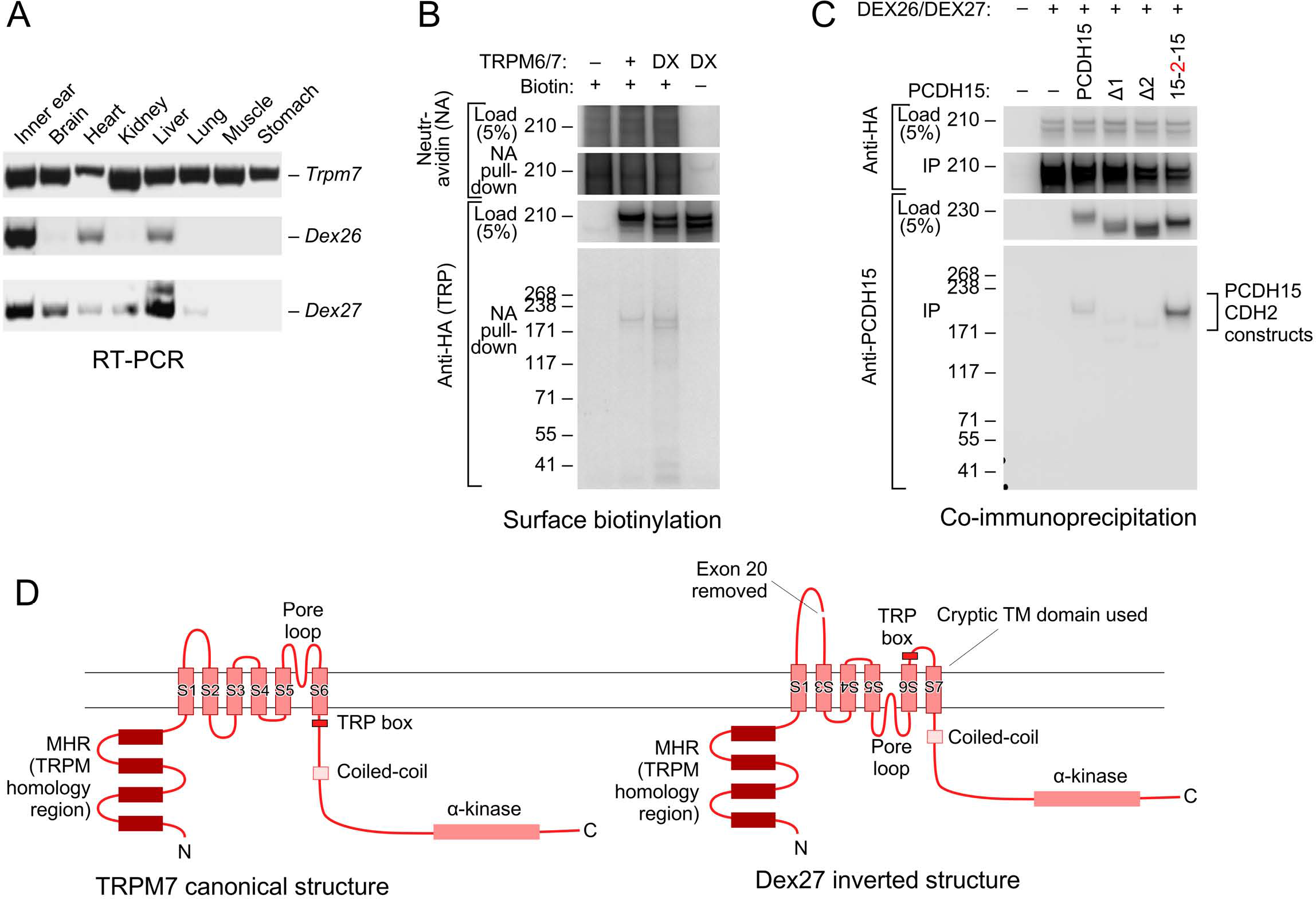
Novel splice forms of *Trpm6* and *Trpm7* interact with PCDH15. ***A***, Tissue expression profiles of mouse *Trpm6, Dex27*, and *Dex26*. Primer sets for *Dex27* and *Dex26* were designed such that they bind to the junctions of *Trpm7* and *Trpm6* exons 19 and 21; signal was only seen if exon 20 was absent. Both *Dex27* and *Dex26* were highly expressed in the ear (first lane on the left). ***B***, Both TRPM6/7 and DEX26/7 proteins traveled to the cell surface. HEK293T cells were transfected with TRPM6/7 or DEX26/7; before harvest, the cells were surface biotinylated. Cell lysates were then immunoprecipitated with neutravidin-agarose, then probed with anti-HA antibody. ***C***, DEX26 and DEX27 interacted with PCDH15 in a similar fashion with TRPM6 and TRPM7. ***D***, Predicted membrane topology for canonical TRPM7 sequence (left) and hypothetical inverted DEX27 sequence (right). The loss of TM2 when exon 20 is spliced out could force inversion of TM3-TM6; a cryptic transmembrane domain (S7 here) is predicted by some topology algorithms to be used, maintaining the N- and C-termini intracellular.

### Mechanotransduction does not require *Trpm7*

These results all suggested the possibility that TRPM6 or TRPM7 mediates mechanotransduction by hair cells. A characteristic of TRPM6 and TRPM7 conductances is that they are blocked by intracellular Mg^2+^ (Nadler et al., 2001; Voets et al., 2004b). Hair-cell transduction was not affected by manipulations of Mg^2+^, however; currents were identical if recorded in the presence of 0 or 3 mM intracellular MgCl_2_ in the internal solution (Fig. 7A).

**Figure 7.**
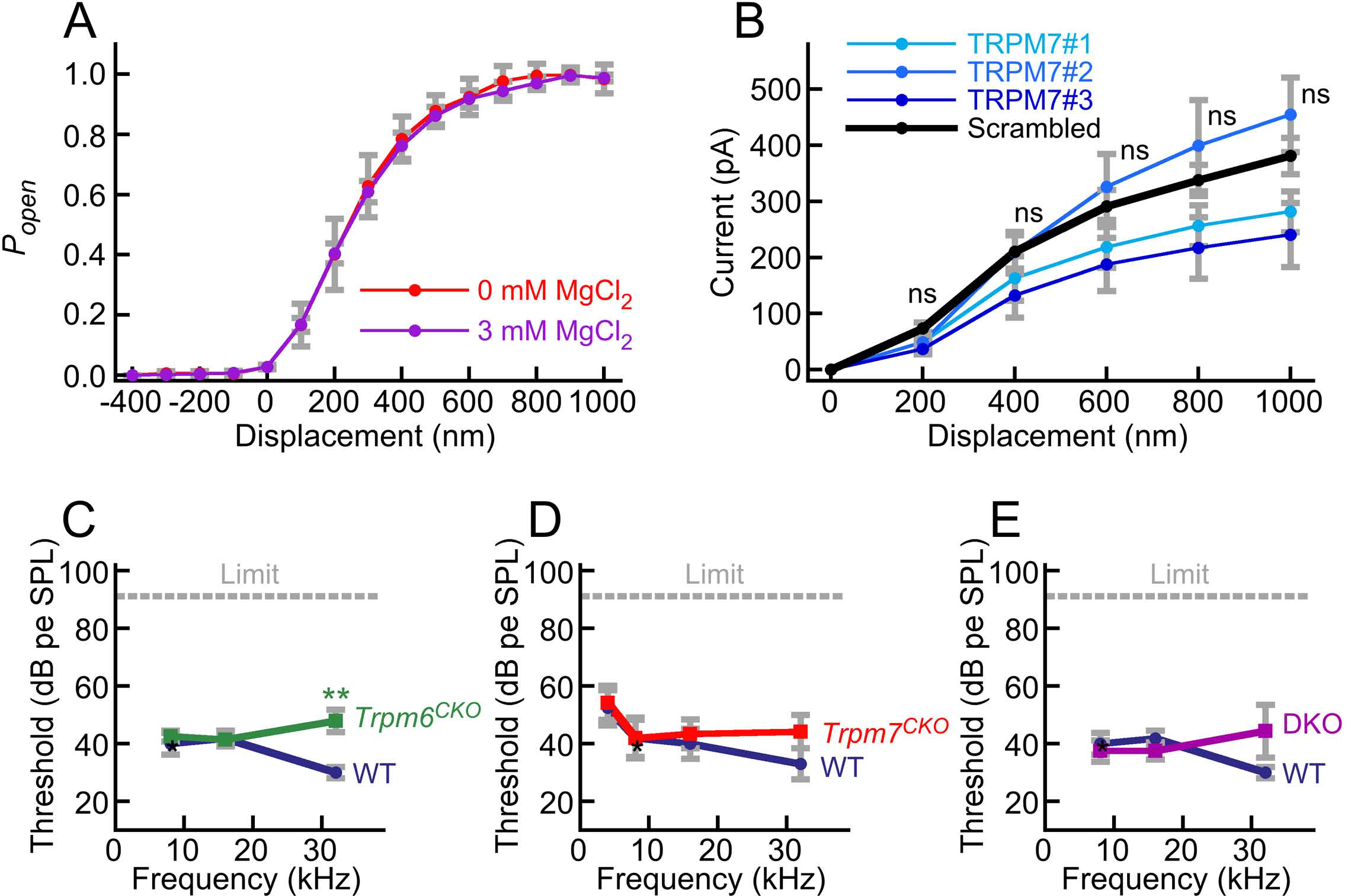
No evidence for contribution of TRPM6 or TRPM7 conductances to hair-cell transduction. ***A***, No effect of internal Mg^2+^. Hair cells were dialyzed with internal solution containing 0 or 3 mM MgCl_2_. No difference in transduction current amplitudes or sensitivity were observed. Mean ± SEM are plotted; n=5 for each. ***B***, shRNAs against *Trpm7* had no effect on mechanotransduction. ns, not significant (for each of the three Trpm7 shRNAs compared to the scrambled shRNA). Mean ± SEM are plotted: TRPM7#1, n=5; TRPM7#2, n=3; TRPM7#3, n=4; scrambled peptide, n=4. ***C***, ABR measurements for *Trpm6*^*CKO*^ mice. Mean ± SEM are plotted: WT, n=11; *Trpm6*^*CKO*^, n=17. **, p<0.01. ***D***, ABR measurements for *Trpm7*^*CKO*^ mice. Mean ± SEM are plotted: WT, n=10; *Trpm*^*CKO*^, n=14. ***E***, ABR measurements for *Trpm6*^*CKO*^; *Trpm7*^*CKO*^ DKO mice. Mean ± SEM are plotted: WT, n=11; *Trpm6*^*CKO*^; *Trpm7*^*CKO*^ DKO, n=8.

We also used a shRNA strategy with injectoporation (Xiong et al., 2014) to interfere with TRPM7 function in hair cells. We confirmed that our shRNAs reduced levels of TRPM7 in HEK293 cells. When cells were transfected with mouse *Trpm7* in the absence of shRNAs, the outward current measured at 200 sec after establishment of the whole-cell recording configuration was 335 ± 22 pA. In cells transfected with mouse *Trpm7* and shRNA TRPM7#1, the current at 200 sec was reduced to 65 ± 26 pA, approximately the same as cells transfected with GFP as a negative control.

We used injectoporation (Xiong et al., 2014) to deliver shRNAs targeting *Trpm7* to cochlear hair cells then measured mechanotransduction using whole-cell recording. Knockdown of *Cdh23* using this paradigm profoundly reduced transduction in transfected cells (Xiong et al., 2014). By contrast, three different *Trpm7* shRNAs each were without significant effect on transduction current amplitude (Fig. 7B).

### *Trpm6*^*CKO*^;*Trpmf*^*CKO*^ double knockout mice have normal hearing

To determine definitively whether TRPM6 or TRPM7 participate in hair-cell mechanotransduction, we produced *Trpm6*^*fl*^ and *Trpm7*^*fl*^ mice (Fig. 2) and expressed CRE recombinase in hair cells to delete essential exons for each gene. Global-null *Trpm6*^*Δ/Δ*^ mice rarely survive past birth due to Mg^2+^ insufficiency (Walder et al., 2009; Woudenberg-Vrenken et al., 2011), and global-null *Trpm7*^*Δ/Δ*^ mice have an embryonic lethality phenotype (Jin et al., 2008). We therefore used the *Atoh1-Cre* transgenic line to restrict *Trpm6* and *Trpm7* deletion to hair cells and a few other cell types (Matei et al., 2005. Pan et al., 2012). Mice with both alleles recombined by CRE are referred to here as *Trpm6*^*CKO*^ or *Trpm7*^*CKO*^.

*Trpm6*^*CKO*^, *Trpm7*^*CKO*^, and *Trpm6*^*CKO*^;*Trpm7*^*CKO*^ (double knockout, or DKO) mice were behaviorally normal, with no indication of gross auditory or vestibular disruption. We assessed auditory function using auditory brainstem response (ABR) measurements. While ABR thresholds at 32 kHz were slightly but significantly elevated for *Trpm6*^*CKO*^ mice (Fig. 7C), thresholds were not statistically significant different for *Trpm7*^*CKO*^ (Fig. 7D) or *Trpm6*^*CKO*^;*Trpm7*^*CKO*^ DkO (Fig. 7E) mice as compared to controls at all frequencies. These results suggest that neither TRPM6 nor TRPM7 is a major contributor the mechanotransduction.

## Discussion

Our experimental results suggest that TRPV5, TRPV6, TRPM6, and TRPM7 are not involved in hair-cell transduction. While each of these TRP channels had plausible initial evidence supporting their involvement, direct experiments—allele-specific inhibition for TRPV5 and TRPV6, and single and double knockouts for TRPM6 and TRPM7—gave no support for a role of any of them in forming the hair cell’s transduction pore. While these results do not provide direct evidence for the participation of TMC1 and TMC2 in forming the pore, the paucity of viable alternative candidates lends indirect support for the two TMCs.

### Inhibition of TRPV5 or TRPV6 does not interfere with hair-cell mechanotransduction

MTS reagents did not inhibit transduction currents measured in hair cells of S556C-*Trpv5* and M527C-*Trpv6* mice. In cell culture, each of these residues can be robustly inhibited from the extracellular side of the membrane (Dodier et al., 2004; Voets et al., 2004a), so the lack of MTS inhibition of transduction in mutant hair cells suggests that TRPV5 and TRPV6 do not contribute to the transduction channel.

To definitively establish the viability of the cysteine substitution/MTS inhibition approach with these channels, future experiments could determine whether Ca^2+^ transport in kidney and intestine, mediated respectively by TRPV5 and TRPV6 (Nijenhuis et al., 2003), is inhibited when these alleles are present and MTS reagents are applied. Like allele-selective inhibition of myosins (Gillespie et al., 1999; Holt et al., 2002), this strategy provides strong evidence for the role of a protein in a biological function; three control conditions (wild-type, wild-type plus inhibitor, mutant) are matched with one experimental condition (mutant plus inhibitor), strengthening the interpretation of an inhibitory effect.

### Elimination of *Trpm6* or *Trpm7* does not interfere with auditory function

Single and double knockouts of *Trpm6* and *Trpm7* had no effect on mouse hearing. While demonstrating conclusively that *Trpm6* and *Trpm7* transcripts are eliminated in hair cells is difficult because of the low level of expression (making in situ mRNA localization challenging) and expression in other cochlear cell types (preventing analysis of whole cochlea to assess transcript loss), recombination by Atoh1-CRE occurs robustly during hair cell development (Matei et al., 2005; Pan et al., 2012). We assessed ABRs at 4-8 weeks of age, and it is very unlikely that *Trpm7* mRNA or TRPM7 protein would be stable for so long after the genes were recombined. Nevertheless, even though there are limitations to our experiments, they nonetheless strongly suggest that TRPM6 and TRPM7 are not part of the transduction channel.

### TRPM6 and TRPM7 bind to PCDH15

Our results suggest that under some circumstances, TRPM6 and TRPM7 can bind to PCDH15, a component of the tip link. Because TRPM6 and TRPM7 are not essential for hearing, these results suggest that TRPM6 and TRPM7 might interact with PCDH15 elsewhere besides the transduction complex. One possibility is that the TRPMs are present in kinocilia, and interact with the PCDH15 molecules that contribute to the kinocilial links (Goodyear et al., 2010). If so, the TRPMs are unlikely to play an essential role in kinocilia-link function, as loss of kinocilial links leads to deafness (Webb et al., 2011) but loss of the TRPMs does not affect hearing. Alternatively, RNA-seq experiments show that PCDH15 is expressed in inner-ear cell types besides hair cells (Burns et al., 2015), and so these channels could form complexes with PCDH15 there.

An alternative view is that PCDH15 or the TRPMs bind membrane proteins promiscuously, which may call into question interactions described based on similar immunoprecipitation experiments (Ramakrishnan et al., 2009; Ramakrishnan et al., 2012; Maeda et al., 2014; Beurg et al., 2015; Cunningham et al., 2017; Erickson et al., 2017). While our experiments were well controlled, including the use of alternative cadherins and TRP channels, reliance simply on cell-culture immunoprecipitation experiments is fraught. Such evidence should always be backed up by alternative experiments that lack some of the ambiguity of in vitro experiments.

### Novel *Trpm6* and *Trpm7* splice forms

We identified splice forms of *Trpm6* and *Trpm7* that each lack exon 20, which encodes most of transmembrane domain 2 in each channel. DEX26 and DEX27 were targeted to the surface of HEK293 cells, and, like the canonical versions of TRPM6 and TRPM7, interacted with PCDH15 via its extracellular domain. Membrane topology programs suggested the possibility that DEX26 and DEX27 have partially inverted transmembrane domain regions, so that the pore of the channel is a re-entrant loop coming from the inside rather than the outside. This topology is speculative at this moment, however, as we have no experimental evidence that the pore is flipped. If this topology is correct, however, it limits the regions of the TRPM proteins that PCDH15 can interact with from the N-terminus through TM2 and from the cryptic TM7 to the C-terminus. Regardless of whether the channel’s pores are inverted, the loss of TM2 will force an unusual topology on this channel, however, and it will be interesting to follow up the role of these splice forms in the organism.

### Implications for identification of the hair cell’s transduction channel

Our results rule out several transduction channel candidates, adding to the list of channels disproven that includes P2RX2, SCNN1A, TRPN1, TRPA1, TRPV4, TRPML3, TRPC3, TRPC5, TRPC6, TRPM1, TRPM2, TRPM3, PKD1, PKD1L3, PKD2, PKD2L1, and PKD2L2 (Rusch and Hummler, 1999; Vollrath et al., 2007; Steigelman et al., 2011; Fettiplace and Kim, 2014; Wu et al., 2016). Because of their essential role in mechanotransduction and their size, the TMCs remain the best candidate for the transduction channel’s pore. Nonetheless, the TMCs still have not been conclusively demonstrated to form ion channels, necessary for showing that a given molecule forms the transduction pore.

The cysteine-substitution/MTS inhibition approach could serve as a useful tool for testing additional transduction channel candidates, although the accurate knowledge of the location of the modified residues from cysteine-scanning mutagenesis (Dodier et al., 2004; Voets et al., 2004a) is essential, as is confirmation of the location of the modified residues in the protein structure (Saotome et al., 2016). Moreover, as modification of cysteines could allosterically affect channel function, perhaps even mediated through protein-protein interactions, the cysteine/MTS approach does not definitively prove that the modified residue is in the tested channel’s pore. Perhaps the most conclusive test for the transduction channel would be to change the ion selectivity of the pore based on knowledge of channel structure, and then show that hair-cell transduction ion selectivity changes in the predicted manner.

## Authors and contributions

CM carried out initial mass spectrometry experiments implicating TRPM6 and TRPM7, and conducted mass spectrometry analysis of immunoprecipitates. HZ demonstrated interaction of TRPM6 and TRPM7 with PCDH15 using immunoprecipitation. ML developed the *Trpv5* and *Trpv6* knock-in mice. WX carried out electrophysiological assays of wild-type mouse hair cells. BP carried out electrophysiological recordings of *Trpv5* and *Trpv6* mice. MA carried out experiments localizing TRPM7. MB assisted in development of *Trpv5* and *Trpv6* knock-in mice, developed genotyping assays for all *Trp* alleles, and conducted ABR measurements. RL was responsible for mouse husbandry and ABRs. AR supervised all TRPV5 and TRPV6 electrophysiology, carried out recordings of *Trpv5* and *Trpv6* mice, and analyzed data. UM supervised TRPV5 and TRPV6 molecular characterization and development of the *Trpm7* floxed allele and analyzed data. PB-G supervised development of *Trpv5* and *Trpv6* knock-in mice, development of *Trpm6* and *Trpm7* knockout mice, and protein interaction experiments; he analyzed data and also wrote the paper with contributions from all authors. All authors agree to be accountable for the content of the work.

## Acknowledgements

We thank Piotr Kazmierczak, Thomas F. Wagner, and Nicolas Grillet for participating in the early phases of this project. We received support from the following core facilities: mouse generation from the Transgenic Mouse Models Core (OHSU), *Trpm6*^*fl*^ embryonic stem cells from KOMP (the Knockout Mouse Project, University of California Davis), and confocal microscopy from the OHSU Advanced Light Microscopy Core @ The Jungers Center.

## References

Adato, A., V. Michel, Y. Kikkawa, J. Reiners, K. N. Alagramam, D. Weil, H. Yonekawa, U. Wolfrum, A. El-Amraoui, and C. Petit (2005). Interactions in the network of Usher syndrome type 1 proteins. Hum Mol Genet. 14, 347–356.

Alagramam, K. N., R. J. Goodyear, R. Geng, D. N. Furness, A. F. van Aken, W. Marcotti, C. J. Kros, and G. P. Richardson (2011). Mutations in protocadherin 15 and cadherin 23 affect tip links and mechanotransduction in mammalian sensory hair cells. PLoS One. 6, e19183.

Asai, Y., J. R. Holt, and G. S. Geleoc (2010). A quantitative analysis of the spatiotemporal pattern of transient receptor potential gene expression in the developing mouse cochlea. J Assoc Res Otolaryngol. 11, 27–37.

Avenarius, M. R., J. F. Krey, R. A. Dumont, C. P. Morgan, C. B. Benson, S. Vijayakumar, C. L. Cunningham, D. I. Scheffer, D. P. Corey, U. Müller, S. M. Jones, and P. G. Barr-Gillespie (2017). Heterodimeric capping protein is required for stereocilia length and width regulation. J. Cell Biol. in press.

Bessac, B. F., and A. Fleig (2007). TRPM7 channel is sensitive to osmotic gradients in human kidney cells. J Physiol. 582, 1073–1086.

Beurg, M., R. Fettiplace, J. H. Nam, and A. J. Ricci (2009). Localization of inner hair cell mechanotransducer channels using high-speed calcium imaging. Nat. Neurosci. 12, 553–558.

Beurg, M., W. Xiong, B. Zhao, U. Müller, and R. Fettiplace (2015). Subunit determination of the conductance of hair-cell mechanotransducer channels. Proc. Natl. Acad. Sci. U. S. A. 112, 1589–1594.

Bianco, S. D., J. B. Peng, H. Takanaga, Y. Suzuki, A. Crescenzi, C. H. Kos, L. Zhuang, M. R. Freeman, C. H. Gouveia, J. Wu, H. Luo, T. Mauro, E. M. Brown, and M. A. Hediger (2006). Marked Disturbance of Calcium Homeostasis in Mice with Targeted Disruption of the Trpv6 Calcium Channel Gene. J Bone Miner Res.

Burns, J. C., M. C. Kelly, M. Hoa, R. J. Morell, and M. W. Kelley (2015). Single-cell RNA-Seq resolves cellular complexity in sensory organs from the neonatal inner ear. Nat Commun. 6, 8557.

Chubanov, V., and T. Gudermann (2014). TRPM6. Handb Exp Pharmacol. 222, 503–520.

Cuajungco, M. P., C. Grimm, and S. Heller (2007). TRP channels as candidates for hearing and balance abnormalities in vertebrates. Biochim. Biophys. Acta. 1772, 1022–1027.

Cunningham, C. L., Z. Wu, A. Jafari, B. Zhao, K. Schrode, S. Harkins-Perry, A. Lauer, and U. Müller (2017). The murine catecholamine methyltransferase mTOMT is essential for mechanotransduction by cochlear hair cells. Elife. 6,

Dodier, Y., U. Banderali, H. Klein, O. Topalak, O. Dafi, M. Simoes, G. Bernatchez, R. Sauve, and L. Parent (2004). Outer pore topology of the ECaC-TRPV5 channel by cysteine scan mutagenesis. J. Biol. Chem. 279, 6853–6862.

Ebrahim, S., M. R. Avenarius, M. Grati, J. F. Krey, A. M. Windsor, A. D. Sousa, A. Ballesteros, R. Cui, B. A. Millis, F. T. Salles, M. A. Baird, M. W. Davidson, S. M. Jones, D. Choi, L. Dong, M. H. Raval, C. M. Yengo, P. G. Barr-Gillespie, and B. Kachar (2016). Stereocilia-staircase spacing is influenced by myosin III motors and their cargos espin-1 and espin-like. Nat Commun. 7, 10833.

Elia, G. (2008). Biotinylation reagents for the study of cell surface proteins. Proteomics. 8, 4012–4024.

Erickson, T., C. P. Morgan, J. Olt, K. Hardy, E. Busch-Nentwich, R. Maeda, R. Clemens, J. F. Krey, A. Nechiporuk, P. G. Barr-Gillespie, W. Marcotti, and T. Nicolson (2017). Integration of Tmc1/2 into the mechanotransduction complex in zebrafish hair cells is regulated by Transmembrane O-methyltransferase (Tomt). Elife. 6,

Fettiplace, R., and K. X. Kim (2014). The physiology of mechanoelectrical transduction channels in hearing. Physiol Rev. 94, 951–986.

Fleig, A., and V. Chubanov (2014). TRPM7. Handb Exp Pharmacol. 222, 521–546.

Gillespie, P. G., S. K. Gillespie, J. A. Mercer, K. Shah, and K. M. Shokat (1999). Engineering of the myosin-Iβ nucleotide-binding pocket to create selective sensitivity to N^6^-modified ADP analogs. J. Biol. Chem. 274, 31373–31381.

Gilliam, J. C., and T. G. Wensel (2011). TRP channel gene expression in the mouse retina. Vision Res. 51, 2440–2452.

Gong, Z., W. Son, Y. D. Chung, J. Kim, D. W. Shin, C. A. McClung, Y. Lee, H. W. Lee, D. J. Chang, B. K. Kaang, H. Cho, U. Oh, J. Hirsh, M. J. Kernan, and C. Kim (2004). Two interdependent TRPV channel subunits, inactive and Nanchung, mediate hearing in Drosophila. J. Neurosci. 24, 9059–9066.

Goodyear, R. J., A. Forge, P. K. Legan, and G. P. Richardson (2010). Asymmetric distribution of cadherin 23 and protocadherin 15 in the kinocilial links of avian sensory hair cells. J. Comp. Neurol. 518, 4288–4297.

Hellwig, N., N. Albrecht, C. Harteneck, G. Schultz, and M. Schaefer (2005). Homo- and heteromeric assembly of TRPV channel subunits. J. Cell Sci. 118, 917–928.

Hoenderop, J. G., and R. J. Bindels (2008). Calciotropic and magnesiotropic TRP channels. Physiology (Bethesda). 23, 32–40.

Hoenderop, J. G., J. P. van Leeuwen, B. C. van der Eerden, F. F. Kersten, A. W. van der Kemp, A. M. Mérillat, J. H. Waarsing, B. C. Rossier, V. Vallon, E. Hummler, and R. J. Bindels (2003). Renal Ca2+ wasting, hyperabsorption, and reduced bone thickness in mice lacking TRPV5. J Clin Invest. 112, 1906–1914.

Holt, J. R., S. K. Gillespie, D. W. Provance, K. Shah, K. M. Shokat, D. P. Corey, J. A. Mercer, and P. G. Gillespie (2002). A chemical-genetic strategy implicates myosin-1c in adaptation by hair cells. Cell. 108, 371–381.

Horwitz, G. C., A. Lelli, G. S. Géléoc, and J. R. Holt (2010). HCN channels are not required for mechanotransduction in sensory hair cells of the mouse inner ear. PLoS One. 5, e8627.

Jin, J., B. N. Desai, B. Navarro, A. Donovan, N. C. Andrews, and D. E. Clapham (2008). Deletion of Trpm7 disrupts embryonic development and thymopoiesis without altering Mg2+ homeostasis. Science. 322, 756–760.

Kawashima, Y., G. S. Geleoc, K. Kurima, V. Labay, A. Lelli, Y. Asai, T. Makishima, D. K. Wu, C. C. Della Santina, J. R. Holt, and A. J. Griffith (2011). Mechanotransduction in mouse inner ear hair cells requires transmembrane channel-like genes. J Clin Invest. 121, 4796–4809.

Kazmierczak, P., H. Sakaguchi, J. Tokita, E. M. Wilson-Kubalek, R. A. Milligan, U. Müller, and B. Kachar (2007). Cadherin 23 and protocadherin 15 interact to form tip-link filaments in sensory hair cells. Nature. 449, 87–91.

Kernan, M. J., (2007). Mechanotransduction and auditory transduction in Drosophila. Pflugers Arch. 454, 703–720.

Krey, J. F., P. A. Wilmarth, J. B. Shin, J. Klimek, N. E. Sherman, E. D. Jeffery, D. Choi, L. L. David, and P. G. Barr-Gillespie (2014). Accurate label-free protein quantitation with high- and low-resolution mass spectrometers. J. Proteome Res. 13, 1034–1044.

Lefevre, G., V. Michel, D. Weil, L. Lepelletier, E. Bizard, U. Wolfrum, J. P. Hardelin, and C. Petit (2008). A core cochlear phenotype in USH1 mouse mutants implicates fibrous links of the hair bundle in its cohesion, orientation and differential growth. Development. 135, 1427–1437.

Lehnert, B. P., A. E. Baker, Q. Gaudry, A. S. Chiang, and R. I. Wilson (2013). Distinct roles of TRP channels in auditory transduction and amplification in Drosophila. Neuron. 77, 115–128.

Maeda, R., K. S. Kindt, W. Mo, C. P. Morgan, T. Erickson, H. Zhao, R. Clemens-Grisham, P. G. Barr-Gillespie, and T. Nicolson (2014). Tip-link protein protocadherin 15 interacts with transmembrane channel-like proteins TMC1 and TMC2. Proc. Natl. Acad. Sci. U. S. A. 111, 12907–12912.

Matei, V., S. Pauley, S. Kaing, D. Rowitch, K. W. Beisel, K. Morris, F. Feng, K. Jones, J. Lee, and B. Fritzsch (2005). Smaller inner ear sensory epithelia in Neurog 1 null mice are related to earlier hair cell cycle exit. Dev Dyn. 234, 633–650.

Nadler, M. J., M. C. Hermosura, K. Inabe, A. L. Perraud, Q. Zhu, A. J. Stokes, T. Kurosaki, J. P. Kinet, R. Penner, A. M. Scharenberg, and A. Fleig (2001). LTRPC7 is a Mg.ATP-regulated divalent cation channel required for cell viability. Nature. 411, 590–595.

Nijenhuis, T., J. G. Hoenderop, B. Nilius, and R. J. Bindels (2003). Patho)physiological implications of the novel epithelial Ca2+ channels TRPV5 and TRPV6. Pflugers Arch. 446, 401–409.

Numata, T., T. Shimizu, and Y. Okada (2007a). Direct mechano-stress sensitivity of TRPM7 channel. Cell Physiol Biochem. 19, 1–8.

Numata, T., T. Shimizu, and Y. Okada (2007b). TRPM7 is a stretch-and swelling-activated cation channel involved in volume regulation in human epithelial cells. Am J Physiol Cell Physiol. 292, C460–7.

Owsianik, G., K. Talavera, T. Voets, and B. Nilius (2006). Permeation and selectivity of TRP channels. Annu Rev Physiol. 68, 685–717.

Pan, B., G. S. Geleoc, Y. Asai, G. C. Horwitz, K. Kurima, K. Ishikawa, Y. Kawashima, A. J. Griffith, and J. R. Holt (2013). TMC1 and TMC2 are components of the mechanotransduction channel in hair cells of the mammalian inner ear. Neuron. 79, 504–515.

Pan, N., I. Jahan, J. Kersigo, J. S. Duncan, B. Kopecky, and B. Fritzsch (2012). A novel Atoh1 “self terminating” mouse model reveals the necessity of proper Atoh1 level and duration for hair cell differentiation and viability. PLoS One. 7, e30358.

Peng, A. W., R. Gnanasambandam, F. Sachs, and A. J. Ricci (2016). Adaptation Independent Modulation of Auditory Hair Cell Mechanotransduction Channel Open Probability Implicates a Role for the Lipid Bilayer. J. Neurosci. 36, 2945–2956.

Powers, R. J., S. Roy, E. Atilgan, W. E. Brownell, S. X. Sun, P. G. Gillespie, and A. A. Spector (2012). Stereocilia membrane deformation: implications for the gating spring and mechanotransduction channel. Biophys. J. 102, 201–210.

Ramakrishnan, N. A., M. J. Drescher, R. L. Barretto, K. W. Beisel, J. S. Hatfield, and D. G. Drescher (2009). Calcium-dependent binding of HCN1 channel protein to hair cell stereociliary tip link protein protocadherin 15 CD3. J. Biol. Chem. 284, 3227–3238.

Ramakrishnan, N. A., M. J. Drescher, K. M. Khan, J. S. Hatfield, and D. G. Drescher (2012). HCN1 and HCN2 Proteins Are Expressed in Cochlear Hair Cells: HCN1 CAN FORM A TERNARY COMPLEX WITH PROTOCADHERIN 15 CD3 AND F-ACTIN-BINDING FILAMIN A OR CAN INTERACT WITH HCN2. J. Biol. Chem. 287, 37628–37646.

Reiners, J., T. Märker, K. Jürgens, B. Reidel, and U. Wolfrum (2005). Photoreceptor expression of the Usher syndrome type 1 protein protocadherin 15 (USH1F) and its interaction with the scaffold protein harmonin (USH1C). Mol Vis. 11, 347–355.

Rusch, A., and E. Hummler (1999). Mechano-electrical transduction in mice lacking the alpha-subunit of the epithelial sodium channel. Hear Res. 131, 170–6.

Saotome, K., A. K. Singh, M. V. Yelshanskaya, and A. I. Sobolevsky (2016). Crystal structure of the epithelial calcium channel TRPV6. Nature. 534, 506–511.

Schwander, M., A. Sczaniecka, N. Grillet, J. S. Bailey, M. Avenarius, H. Najmabadi, B. M. Steffy, G. C. Federe, E. A. Lagler, R. Banan, R. Hice, L. Grabowski-Boase, E. M. Keithley, A. F. Ryan, G. D. Housley, T. Wiltshire, R. J. Smith, L. M. Tarantino, and U. Müller (2007). A forward genetics screen in mice identifies recessive deafness traits and reveals that pejvakin is essential for outer hair cell function. J. Neurosci. 27, 2163–2175.

Siemens, J., C. Lillo, R. A. Dumont, A. Reynolds, D. S. Williams, P. G. Gillespie, and U. Müller (2004). Cadherin 23 is a component of the tip link in hair-cell stereocilia. Nature. 428, 950–955.

Sollner, C., G. J. Rauch, J. Siemens, R. Geisler, S. C. Schuster, U. Müller, and T. Nicolson (2004). Mutations in cadherin 23 affect tip links in zebrafish sensory hair cells. Nature. 428, 955–959.

Steigelman, K. A., A. Lelli, X. Wu, J. Gao, S. Lin, K. Piontek, C. Wodarczyk, A. Boletta, H. Kim, F. Qian, G. Germino, G. S. Géléoc, J. R. Holt, and J. Zuo (2011). Polycystin-1 is required forstereocilia structure but not for mechanotransduction in inner ear hair cells. J. Neurosci. 31, 12241–12250.

Venkatachalam, K., and C. Montell (2007). TRP channels. Annu Rev Biochem. 76, 387–417.

Voets, T., A. Janssens, G. Droogmans, and B. Nilius (2004a). Outer pore architecture of a Ca^2+^-selective TRP channel. J. Biol. Chem. 279, 15223–15230.

Voets, T., B. Nilius, S. Hoefs, A. W. van der Kemp, G. Droogmans, R. J. Bindels, and J. G. Hoenderop (2004b). TRPM6 forms the Mg2+ influx channel involved in intestinal and renal Mg2+ absorption. J. Biol. Chem. 279, 19–25.

Vollrath, M. A., K. Y. Kwan, and D. P. Corey (2007). The micromachinery of mechanotransduction in hair cells. Annu Rev Neurosci. 30, 339–365.

Walder, R. Y., B. Yang, J. B. Stokes, P. A. Kirby, X. Cao, P. Shi, C. C. Searby, R. F. Husted, and V. C. Sheffield (2009). Mice defective in Trpm6 show embryonic mortality and neural tube defects. Hum Mol Genet. 18, 4367–4375.

Webb, S. W., N. Grillet, L. R. Andrade, W. Xiong, L. Swarthout, C. C. Della Santina, B. Kachar, and U. Muller (2011). Regulation of PCDH15 function in mechanosensory hair cells by alternative splicing of the cytoplasmic domain. Development. 138, 1607–1617.

Wei, C., X. Wang, M. Chen, K. Ouyang, L. S. Song, and H. Cheng (2009). Calcium flickers steer cell migration. Nature. 457, 901–905.

Wilmarth, P. A., M. A. Riviere, and L. L. David (2009). Techniques for accurate protein identification in shotgun proteomic studies of human, mouse, bovine, and chicken lenses. J. Ocul. Biol. Dis. Infor. 2, 223–234.

Woudenberg-Vrenken, T. E., A. Sukinta, A. W. van der Kemp, R. J. Bindels, and J. G. Hoenderop (2011). Transient receptor potential melastatin 6 knockout mice are lethal whereas heterozygous deletion results in mild hypomagnesemia. Nephron Physiol. 117, p11–9.

Wu, X., A. A. Indzhykulian, P. D. Niksch, R. M. Webber, M. Garcia-Gonzalez, T. Watnick, J. Zhou, M. A. Vollrath, and D. P. Corey (2016). Hair-Cell Mechanotransduction Persists in TRP Channel Knockout Mice. PLoS One. 11, e0155577.

Xiao, E., H. Q. Yang, Y. H. Gan, D. H. Duan, L. H. He, Y. Guo, S. Q. Wang, and Y. Zhang (2015). Brief reports: TRPM7 Senses mechanical stimulation inducing osteogenesis in human bone marrow mesenchymal stem cells. Stem Cells. 33, 615–621.

Xiong, W., N. Grillet, H. M. Elledge, T. F. Wagner, B. Zhao, K. R. Johnson, P. Kazmierczak, and U. Muller (2012). TMHS is an integral component of the mechanotransduction machinery of cochlear hair cells. Cell. 151, 1283–1295.

Xiong, W., T. Wagner, L. Yan, N. Grillet, and U. Muller (2014). Using injectoporation to deliver genes to mechanosensory hair cells. Nat Protoc. 9, 2438–2449.

Zanini, D., and M. C. Göpfert (2014). TRPs in hearing. Handb Exp Pharmacol. 223, 899–916.

Zhang, W., Z. Yan, L. Y. Jan, and Y. N. Jan (2013). Sound response mediated by the TRP channels NOMPC, NANCHUNG, and INACTIVE in chordotonal organs of Drosophila larvae. Proc. Natl. Acad. Sci. U. S. A. 110, 13612–13617.

Zhao, B., Z. Wu, N. Grillet, L. Yan, W. Xiong, S. Harkins-Perry, and U. Muller (2014). TMIE is an essential component of the mechanotransduction machinery of cochlear hair cells. Neuron. 84, 954–967.

Zhao, H., M. R. Avenarius, and P. G. Gillespie (2012). Improved biolistic transfection of hair cells. PLoS One. 7, e46765.

